# Aerosolizable plasmid DNA dry powders engineered by thin-film freezing

**DOI:** 10.1101/2022.10.03.510625

**Authors:** Haiyue Xu, Chaeho Moon, Sawittree Sahakijpijarn, Huy M. Dao, Riyad F. Alzhrani, Jie-liang Wang, Robert O. Williams, Zhengrong Cui

## Abstract

This study was designed to test the feasibility of using thin-film freezing (TFF) to prepare aerosolizable dry powders of plasmid DNA (pDNA) for pulmonary delivery. Dry powders of pDNA formulated with mannitol/leucine (70/30, w/w) at various of drug loadings, solid contents, and solvents were prepared using TFF, their aerosol properties (i.e., mass median aerodynamic diameter (MMAD) and fine particle fraction (FPF)) determined, and selected powders were used for further characterization. Of the nine dry powders prepared, their MMAD values were about 1-2 mm, with FPF values (delivered) of 40-80%. The aerosol properties of the powders were inversely correlated with the pDNA loading and the solid content in the pDNA solution before thin-film freezing. Powders prepared with Tris-EDTA (TE) buffer or cosolvents (i.e., 1,4 dioxane or t-butanol in water), instead of water, showed slightly reduced aerosol properties. Ultimately, powders prepared with pDNA loading at 5% (w/w), 0.25% of solid content, with or without TE were selected for further characterization due to their overall good aerosol performance. The pDNA powders exhibited a porous matrix, crystalline structure, with a moisture content of <2% (w/w). Agarose gel electrophoresis confirmed the chemical integrity of the pDNA after it was subjected to TFF and after the TFF powder was actuated. A cell transfection study confirmed the activity of the pDNA after it was subjected to TFF. In conclusion, it is feasible to use TFF to produce aerosolizable pDNA dry powder for pulmonary delivery, while preserving the integrity and activity of the pDNA.

## INTRODUCTION

The lung is an appealing target for administering nucleic acid-based products such as plasmid DNA (pDNA) for vaccination by oral inhalation, due to the non-invasive nature of pulmonary delivery and the vast and highly vascularized surface area of the lung [1, 2]. Moreover, pulmonary delivery of pDNA holds potential in treating various lung diseases caused by gene mutations, such as cystic fibrosis, alpha-1 antitrypsin deficiency, acute illnesses such as acute transplant rejection, as well as lung cancer [3-6]. However, effective pulmonary administration of pDNA has been challenging. Only aerosol droplets or particles with aerodynamic diameters in the range of 1 to 5 μm can reach and contact bronchial and alveolar epithelial cells upon inhalation [7]. However, pulmonary delivery of naked pDNA is challenging using traditional nebulizers, including jet and ultrasonic nebulizers, due to the shear and cavitational stresses associated with nebulization [8-10]. The collapse of air bubbles creates shock waves that can damage the pDNA, with their tertiary structure changing from supercoiled to open circular (i.e., nicking form) and/or fragmented configurations (e.g., linear form or even small fragments) [8-10]. The sensitivity of pDNA to shearing is strongly correlated with plasmid length; plasmids of above 5 kb are significantly more sensitive to shear-induced degradation than smaller plasmids [8, 11]. Unfortunately, the damage caused by shearing can be consequential, causing the transfection efficiency of pDNA after nebulization to decrease to 10% [12].

Pulmonary delivery of pDNA in a dry powder form by oral inhalation using a dry powder inhaler (DPI) is an alternative method to nebulization of pDNA in liquid. However, dry powder engineering technologies such as spray drying and spray freeze-drying that can generate powders with good aerosol properties for lung delivery also involve atomization of liquid by spraying, and the shear stress associated with the atomization has proven damaging to pDNA [13, 14]. For example, Kuo and Hwang (2003) reported that spray freeze-drying adversely affects the tertiary structure of pDNA, resulting in linear form of DNA even in the presence of protective agents such as sucrose, trehalose, or mannitol. The similar pDNA damage was also reported when the pDNA was subjected to spray drying [15, 16].

Thin-film freezing (TFF) is an ultra-rapid freezing technology, which employs a cryogenically cooled solid surface to freeze samples. Small droplets of liquid (∼2 mm in diameter) are dropped from above the surface. Upon impact, the droplet spreads and is then frozen in 70-3000 milliseconds. Drying of the frozen thin films by lyophilization generates powders often with desirable aerosol properties, due to their low-density, large specific surface area, and brittle matrix nature [17-19]. TFF technology has been used to prepare aerosolizable dry powders of small molecule drugs for pulmonary delivery, such as tacrolimus, remdesivir, voriconazole, and niclosamide [20-25]. Recently, it has also been applied to large molecules such as monoclonal antibodies and enzymes and particulates such as liposomes and small interfering RNA-solid lipid nanoparticles [26-32]. Due to the low shear stress associated with TFF, using TFF to prepare dry powders minimizes the detrimental effect from shear stress to large molecules such as monoclonal antibodies and enzymes, as compared to spray freeze-drying and spray drying [33].

In this study, two plasmids that encode β-galactosidase (β-gal) or green fluorescent protein (GFP) were employed to study the feasibility of applying TFF to prepare aerosolizable dry powders of plasmids. The pDNA powders were prepared with mannitol and leucine (70:30, w/w) as excipients as powders prepared with this specific excipient composition generally have good aerosol properties [34]. The effect of Tris-EDTA (TE) buffer and co-solvents including 1,4-dioxane or *Tert*-butanol on the aerosol performance properties of the resultant dry powders were also evaluated. Finally, the integrity of pDNA after being subjected to thin-film freezing (TFF) and actuation using a DPI device was tested using agarose gel electrophoresis and/or transfection of A549 human lung epithelial-like cells in culture.

## MATERIALS AND METHODS

### Materials

The β-galactosidase gene-encoding pDNA pCMV-β was from the American Type Culture Collection (ATCC, Manassas, VA). It was constructed based on pUC19 plasmid with a AMP^r^ gene, capable of expressing *E*.*coli* β-Gal under the control of different viral promoters in mammalian cells [35]. The GFP expressing plasmid pVectOZ-GFP was from OZbiosciences (San Diego, CA), which contains a modified human cytomegalovirus (CMV) promoter and a KAN^r^ gene. DH5α competent cells, LB broth, cell extraction buffer and Lipofectamine 3000 reagent were from Invitrogen (Carlsbad, CA). Plasmid Midiprep kit and Maxi kit were from QIAGEN (Valencia, CA). The 1,4-dioxane, *tert*-butanol, TE buffer, and ampicillin were from Fisher Scientific (Fair Lawn, NJ). Restriction digestion enzymes *Hind* III, *Eco*RI, and *Bam*HI were from New England Biolabs (Ipswich, MA). Agarose powder was from Amresco (Atlanta, GA). Polysorbate 20, lactose monohydrate, and methanol anhydrate were from Sigma-Aldrich (St. Louis, MO). GeneRuler 1 kb Plus DNA ladder and Quant-iT™ PicoGreen™ dsDNA Assay Kit were from Thermo Scientific (Waltham, MA). Size #3 hydroxypropyl methylcellulose Quali-V-I capsules were from Qualicaps (Whitsett, NC). GFP ELISA kit was from Abcam (Waltham, MA). A549 cells from ATCC (Manassas, VA) were grown in Roswell Park Memorial Institute (RPMI) medium supplemented with 10% (v/v) fetal bovine serum (FBS) and penicillin–streptomycin at final concentrations of 100 U/mL and 100 μg/mL, respectively. The cell culture reagents including Trypsin-EDTA used in the subculture were all from Gibco (Grand Island, NY).

### Plasmid Preparation

The pCMV-β or pVectOZ-GFP was transformed into *E. coli* DH5α under selective growth conditions and then amplified and purified using a plasmid Midiprep kit. Large scale plasmid preparation was performed using plasmid Maxi kit. The plasmid concentration was evaluated using a Nanodrop 2000 from Thermo Scientific.

### Preparation of Plasmid DNA Dry Powders Using Thin-Film Freezing

To screen for the best dry powder formulation for oral inhalation into the lung, pCMV-β or pVectOZ-GFP with mannitol and leucine (70:30, w/w) as excipients were dissolved in either water, TE buffer, 1,4-dioxane/water (10/90, *v*/*v*), or *Tert*-butanol/water (40/60, *v*/*v*) at various solid contents and plasmid loading levels as shown in Table 1. The formulations were temporarily stored in a refrigerator at ∼4°C before being applied to the TFF process.

**Table 1.**
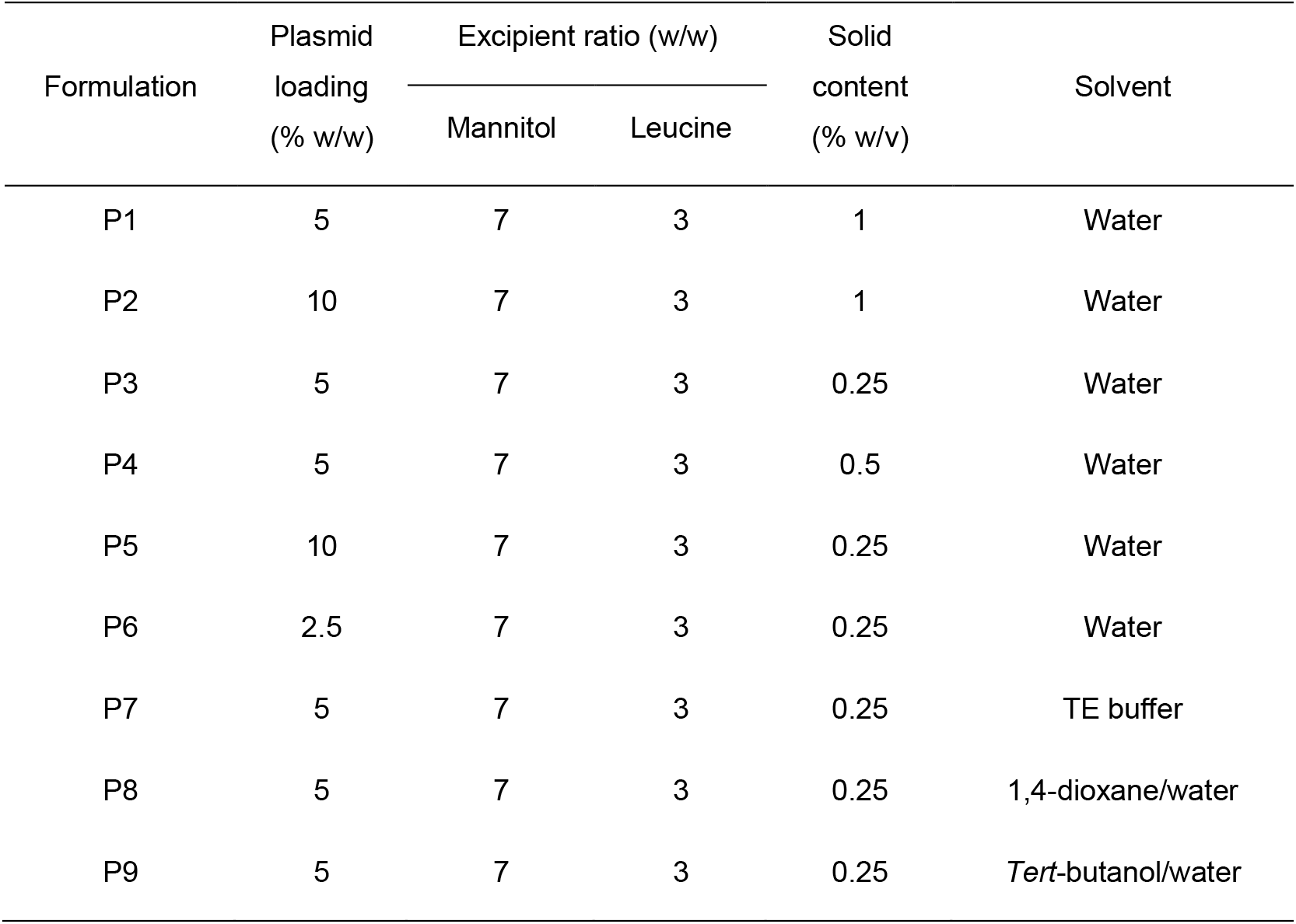
List of plasmid compositions.

| Formulation | Plasmid loading<br>(% w/w) | Excipient ratio (w/w) |  | Solid content<br>(% w/v) | Solvent |
| --- | --- | --- | --- | --- | --- |
|  |  | Mannitol | Leucine |  |  |
| P1 | 5 | 7 | 3 | 1 | Water |
| P2 | 10 | 7 | 3 | 1 | Water |
| P3 | 5 | 7 | 3 | 0.25 | Water |
| P4 | 5 | 7 | 3 | 0.5 | Water |
| P5 | 10 | 7 | 3 | 0.25 | Water |
| P6 | 2.5 | 7 | 3 | 0.25 | Water |
| P7 | 5 | 7 | 3 | 0.25 | TE buffer |
| P8 | 5 | 7 | 3 | 0.25 | 1,4-dioxane/water |
| P9 | 5 | 7 | 3 | 0.25 | *Tert*-butanol/water |

The TFF process and lyophilization were done as previously described [32, 34, 36, 37]. Briefly, 0.25 mL of sample was dropped through a 21-gauge syringe dropwise onto a rotating cryogenically cooled stainless-steel surface (-80°C). To form frozen thin films, the speed at which the surface of the drum rotated was controlled at 5–7 rpm to avoid the overlap of droplets. The frozen thin films were removed using a steel blade and collected in liquid nitrogen in a glass vial. The glass vial was capped with a rubber stopper with half open and transferred into a −80°C freezer for a temporary storage, and then transferred to a VirTis Advantage bench top tray lyophilizer with stopper re-cap function (The VirTis Company, Inc. Gardiner, NY). Lyophilization was performed over 60 h at pressures no more than 100 mTorr, while the shelf temperature was gradually ramped from −40°C to 25°C. The lyophilization cycle is shown in Table 2.

**Table 2.** Lyophilization cycle used to dry the thin-film frozen plasmids.

| Lyophilization Stage | Parameters |
| --- | --- |
| Loading/Freezing temp | -40°C |
| Primary drying temp | -40°C |
| Primary drying time | 20 h |
| Ramp to secondary drying | 20 h |
| Secondary drying temp | +25°C |
| Secondary drying time | 20 h |

### In Vitro Aerosol Performance Evaluation

The aerosol performance properties of the thin-film freeze-dried (TFFD) plasmid powders were determined as previously described [32, 34, 36, 37]. Briefly, a Next Generation Pharmaceutical Impactor (NGI) (MSP Corp, Shoreview, MN) connected to a High-Capacity Pump (model HCP5, Copley Scientific, Nottingham, UK) and a Critical Flow Controller (model TPK 2000, Copley Scientific, Nottingham, UK) was adopted to assess the aerosol performance. To avoid bounce of emitted particles across NGI collection plates, the plates were precoated with 1.5%, *w*/*v*, polysorbate 20 in methanol and dried in air before use. Plasmid DNA powder (2–3 mg) was loaded into a Size #3 capsule, and the capsule was loaded into a high-resistance Plastiape® RS00 inhaler (Plastiape S.p.A, Osnago, Italy) attached to a United States Pharmacopeia (USP) induction port (Copley Scientific, Nottingham, UK). The powder was dispersed to the NGI at the flow rate of 60 L/min for 4 s per actuation, providing a 4 kPa pressure drop across the device. Then, the powders in the capsule, inhaler, adapter, induction port, stages 1–7, and the micro-orifice collector (MOC) were collected by dissolving them with water, and the amount of plasmid DNA in samples was quantified using a PicoGreen™ dsDNA Assay Kit following the manufacturer’s instruction.

The Copley Inhaler Testing Data Analysis Software (CITDAS) Version 3.10 (Copley Scientific, Nottingham, UK) was used to calculate the MMAD, the geometric standard deviation (GSD), and the FPF values. The FPF of recovered dose was calculated as the total amount of plasmid collected with an aerodynamic diameter below 5 μm as a percentage of the total amount of plasmid collected. The FPF of delivered dose was calculated as the total amount of plasmids collected with an aerodynamic diameter below 5 μm as a percentage of the total amount plasmids deposited on the adapter, the induction port, stages 1–7 and MOC of the NGI device.

### Scanning Electron Microscopy (SEM)

The morphology of powder was examined using a Zeiss Supra 40C scanning electron microscope (Carl Zeiss, Heidenheim an der Brenz, Germany) in the Institute for Cell and Molecular Biology Microscopy and Imaging Facility at The University of Texas at Austin. A small amount of bulk powder (i.e., a flake of TFF powder) was deposited on the specimen stub using a double-stick carbon tape. A sputter was used to coat the sample with 15 nm of 60/40 of Pd/Pt before capturing images.

### Moisture Content Measurement

TFF powder (10 mg, n = 3) was diluted into CombiMethanol solvent from Aquastar (Darmstadt, Germany), and the moisture was measured and determined using a Mettler Toledo V20 volumetric Karl Fischer (KF) titrator (Columbus, OH).

### X-ray Powder Diffraction (XRPD)

A Rigaku Miniflex 600 II (Rigaku, Tokyo, Japan) equipped with primary monochromated radiation (Cu K radiation source, λ = 1.54056 Å) was adopted for XRPD study. Plasmid pCMV-β powder sample was loaded onto the sample holder and then analyzed in continuous mode. The operating conditions of accelerating voltage of 40 kV was at 15 mA, step size of 0.02° over a 2θ range of 5-40°, scan speed of 1°/min, and dwell time of 2 s as previously described [26].

### Modulated Differential Scanning Calorimetry (mDSC)

Plasmid pCMV-β powder (3-5 mg) was accurately weighed and loaded into Tzero aluminum hermetic crucibles. A puncture was made in the top lid, right before the DSC measurement. A Model Q20 (TA Instruments, New Castle, DE) differential scanning calorimeter equipped with a refrigerated cooling system (RCS40, TA Instruments, New Castle, DE) was used. In the measurement process, samples were first cooled down to −40°C at a rate of 10oC/min and then ramped up from −40 to 300°C at a rate of 5°C/min. The rate of dry nitrogen gas flow was set as 50 mL/min. The scans were performed with a modulation period of 60 s and a modulated amplitude of 1°C. A TA Instruments Trios v.5.1.1.46572 software was used to analyze the data.

### Restriction Digestion and Agarose Gel Electrophoresis

Plasmid pCMV-β or pVectOZ-GFP was formulated into formulation P7 (see Table 1) and thin-film freeze-dried. The pCMV-β powder was reconstituted and then digested with *EcoR* I alone or *Hind* III and *EcoR* I for 2 h at 37°C. The pVectOZ-GFP plasmid dry powder was reconstituted and then digested with *Hind* III alone or both *Hind* III and *BamH* I for 2 h at 37°C. Final restriction digestion products were applied to agarose gel (0.8%) for electrophoresis. Controls include plasmid alone or plasmid in formulation P7 without being subjected to TFFD, both restriction-digested before electrophoresis.

### In Vitro Transfection Study with A549 Cells

A549 cells were seeded in a 24-well plate (2.1 × 10^5^/well) and incubated at 37°C, 5% CO_2_. An 80% of confluence was reached after 24 h. To prepare the cell transfection reagents, pVectOZ-GFP in formulation P7 before and after being subjected to TFF (i.e., dry powder reconstituted with water) was mixed with Lipofectamine 3000 following the manufacturer’s instruction. Cells were treated with the reagent (amount fixed) mixed with 100, 250, 500, 1000 and 2500 ng of pDNA/well in formulation P7 before being subjected to TFF to determine the optimum dose for cell transfection, and 500 ng of pDNA/well was selected to compare GFP expression by pVectOZ-GFP before and after being subjected to TFF. The cell medium was changed to fresh medium after 20 h, and cells were then incubated for another 24 h. Finally, cells were washed with 1 × PBS, harvested, and suspended in a cell lysis buffer for 30 min. The supernatant was collected and measured with a GFP ELISA kit following the manufacturer’s instruction. Controls include Lipofectamine 3000 alone and cells left untreated.

### Spray Freezing and Spray Freeze-Drying

Plasmid pCMV-β in formulation 7 (Table 1) were spray-atomized using a 2-fluid nozzle (BUCHI Corporation, DE, New Castle) at different air-flow rates (20, 15, 5 or 2.5 L/min) with high purity nitrogen. The air-flow rate was controlled via a critical flow controller (TPK 2000, Copley Scientific, Nottingham, UK) positioned at the nozzle input. The resultant droplets with a diameter ranging from 30-50 µm were atomized into a liquid nitrogen containing Erlenmeyer flask submerged in liquid nitrogen. The resultant frozen material containing flask was placed in a -80°C freezer to allow the liquid nitrogen to evaporate and thawed at 4°C for agarose gel electrophoresis. For spray freeze-drying, the spray-frozen material was dried in a bench lyophilizer following the drying procedures shown in Table 2.

## RESULTS AND DISCUSSION

### In Vitro Aerosol Performance

The aerosol performance properties of the TFF pDNA powders are shown in Fig. 1 and Table 3. Overall, all nine powders showed good aerosol properties. However, it is also clear that dry powders prepared with lower solid contents showed better aerosol performance. For example, the FPF_<5 μm_ values (of the delivered dose) of the plasmid formulations prepared with 1.0, 0.5 and 0.25%, w/v, of solid content (i.e., P1, P4 and P3) were 57.26 ± 3.19%, 56.63 ± 3.82% and 72.32 ± 0.41%, respectively, and the MMAD values of these powders were 1.58 ± 0.07 μm, 1.77 ± 0.22 μm and 1.44 ± 0.16 μm, respectively (Table 3 and Fig. 2A). As to the effect of the plasmid loading (i.e., plasmid weight vs. total weight) on the aerosol performance, lower plasmid loading showed better aerosol performance. For example, the FPF_<5 μm_ values (of the delivered dose) of plasmid formulations prepared with 10.0, 5.0 and 2.5%, w/w, of plasmid (i.e., P5, P3 and P6, respectively) were 46.16 ± 4.44%, 72.32 ± 0.41% and 80.39 ± 3.23%, respectively, and the MMAD values of these powders were 1.69 ± 0.30 μm, 1.44 ± 0.16 μm and 1.27 ± 0.40 μm, respectively (Table 3 and Fig. 2B). This was likely due to the polymeric nature of plasmid DNA. At higher plasmid loading (e.g., 10% vs. 2.5%), stronger intermolecular interactions may have made the powders more difficult to break when they were actuated from the DPI device.

**Table 3.**
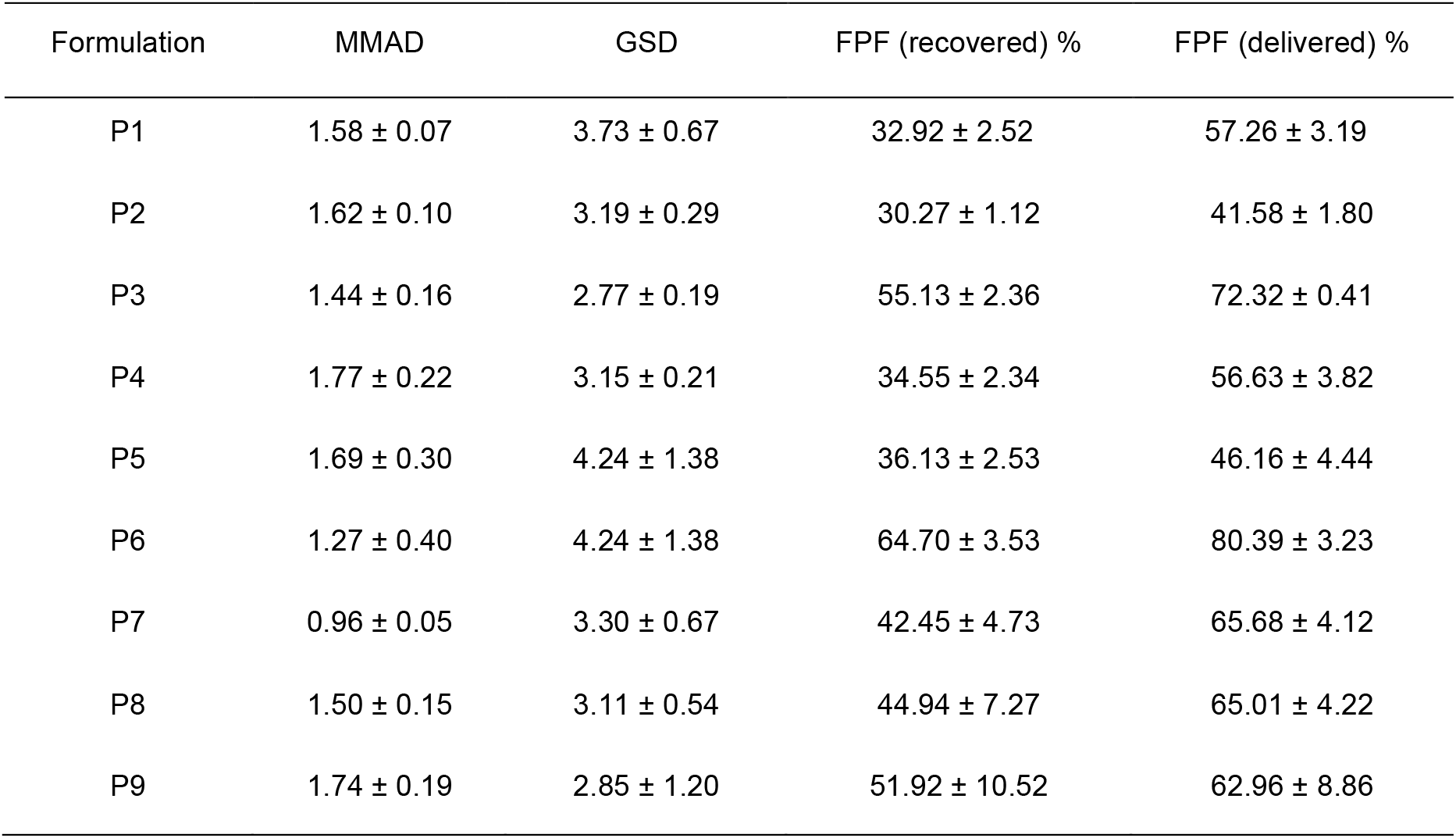
Aerosol performance properties of thin-film freeze-dried pCMV-β powders. Data are mean ± S.D. (*n* = 3) (MMAD, mass median aerodynamic diameter; GSD, geometric standard deviation; FPF, fine particle fraction).

**Figure 1.**
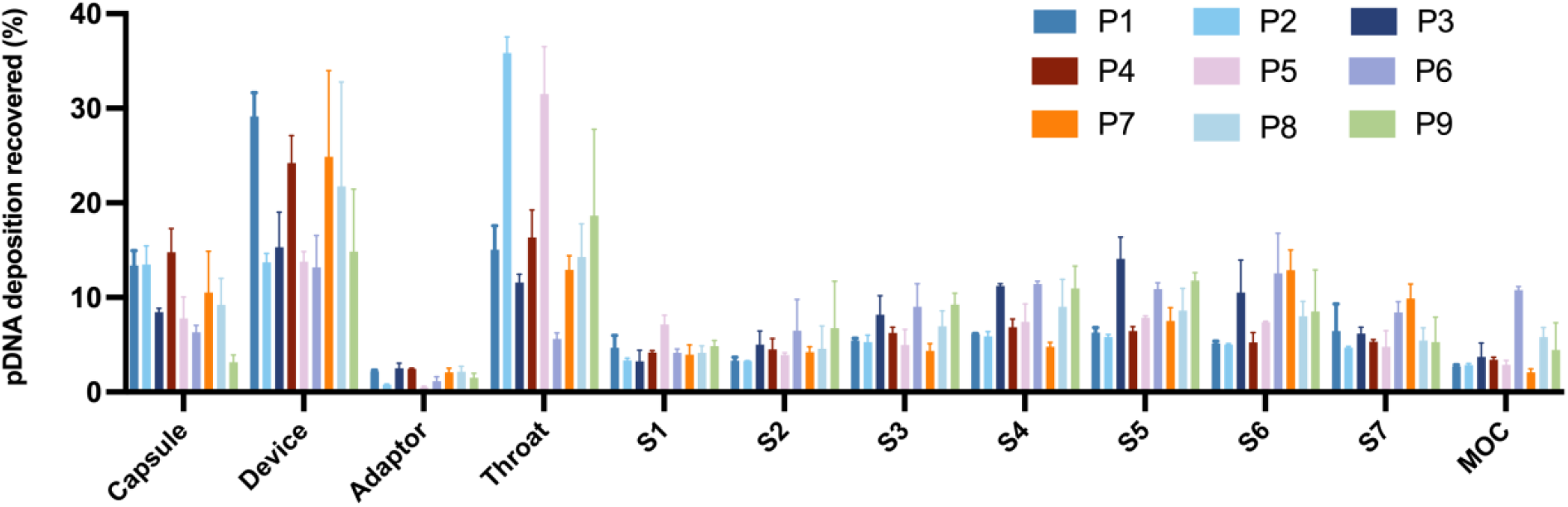
Deposition profiles of TFF plasmid DNA powders in various stages after the powders were applied to NGI using a Plastiape® RS00 high-resistance DPI at a flow rate of 60 L/min. Data are mean ± S.D. (*n* = 3)

**Figure 2.**
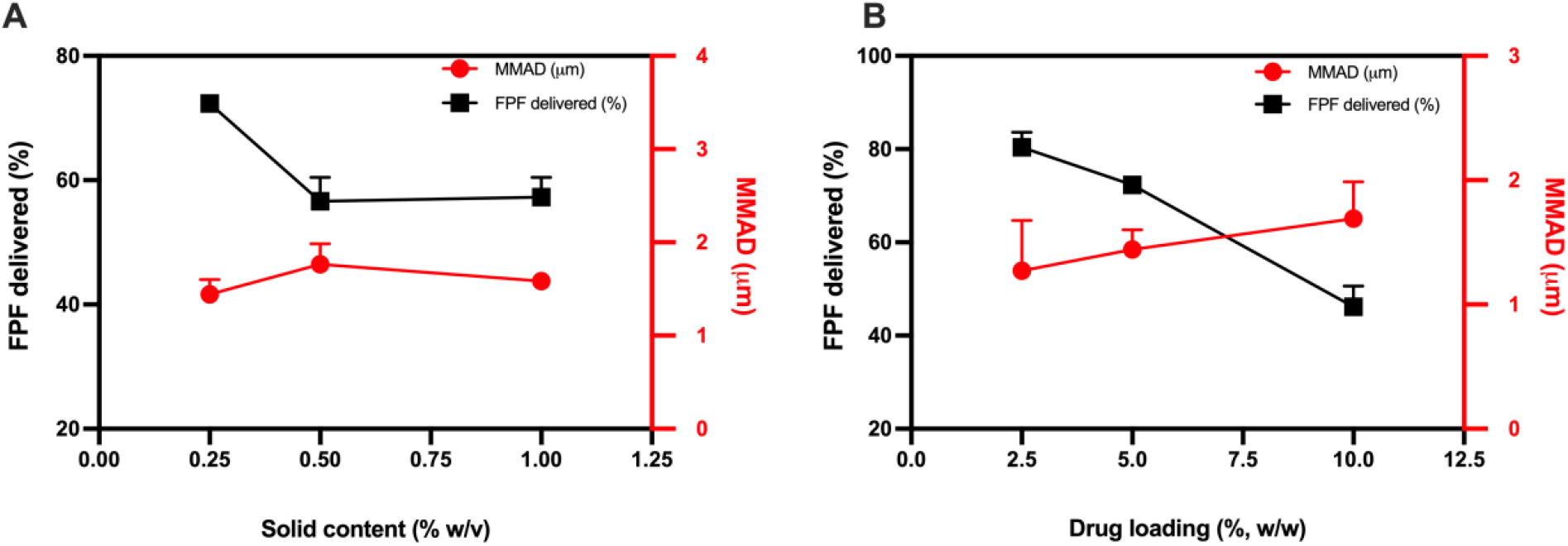
Correlation of solid content (A) and pDNA loading (B) with aerosol performance of TFF plasmid DNA powders. Data are mean ± S.D. (*n* = 3)

The effects of co-solvent and TE buffer on the aerosol performance were also investigated. TE buffer was included with the intention to protect pDNA from DNase digestion as the EDTA in the TE buffer is a chelator of divalent cations such as Mg^2+^, which are required for the DNase activity [38]. The 1,4-dioxane and *Tert*-butanol were used to prepare cosolvents because data from our previous studies showed that they help increase the solubility of certain molecules in water and improve the aerosol properties of the resultant TFF powders [37]. Overall, including TE buffer, 1,4-dioxane, or *Tert*-butanol in the solvent did not improve the FPF_<5 μm_ (Fig. 1, Table 3). It appeared that including the TE buffer in the solvent led to a slight decrease in the aerosol performance properties of the resultant dry powder (i.e., P3 vs. P7, Fig. 1 and Table 3). If the stability of the pDNA during long term storage needs improvement, then the TE buffer or EDTA alone may be included in the powder. The slightly negative effect of the 1,4-dioxane/water or t-butanol/water cosolvents on the aerosol performance of the resultant pDNA powders may be attributed to the highly water-soluble nature of the pDNA. Ultimately, formulations P3 and P7 were chosen for additional characterization because they both have desirable aerosol properties and contained a relatively high amount of the pDNA (i.e., 5% pDNA loading).

### Physical Characteristics of Thin-Film Freeze-Dried Plasmid DNA Powders

The moisture content in the pDNA powder formulation P3 was 1.59 ± 0.12% (w/w). SEM images revealed that the pDNA powder formulation P3 contained nanostructured aggregates (Fig. 3A-B), with highly porous matrix structure (Fig. 3C), which explains the good aerosol performance properties of the powder as shown in Fig. 1 and Table 3.

**Figure 3.**
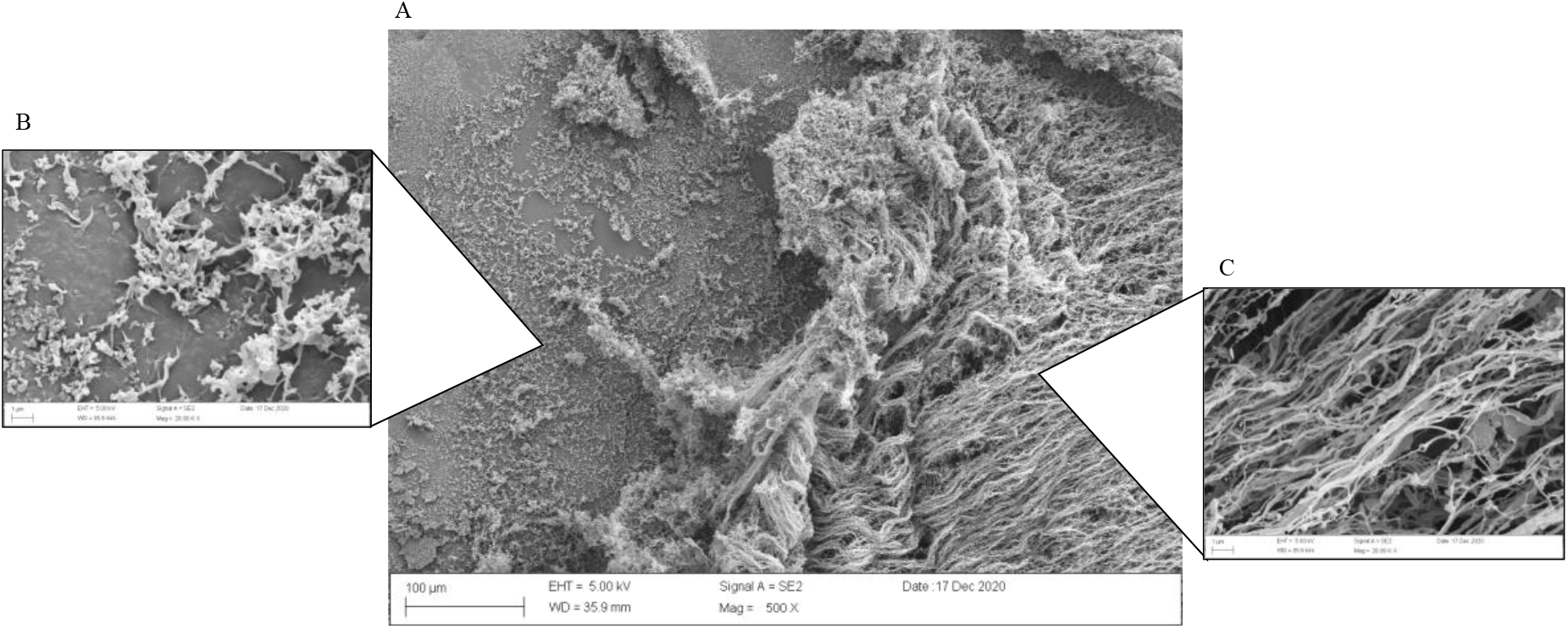
Representative SEM images of a TFF pCMV-β plasmid powder (i.e., formulation P3).

XRPD and DSC were carried out to analyze pDNA powder formulation 7. XRPD diffractogram (Fig. 4) showed that TE salt was in crystalline form after the TFF process as several sharp peaks were observed in the TFF neat TE salts (e.g., 10.8, 14, 15.2, 18.2, 20.2, 21.5, 22.5, 23.5, 26, 27, 27.5, 31, 32.2, 33, 34, and 39.2-degree two-theta) and TFF pDNA P7 formulation (e.g., 10.8 and 15.2 degree two-theta). Sharp peaks of mannitol were observed in TFF neat mannitol (e.g., 9.5, 13.5, 14.5 17, 18.5, 20.2, 21, 22, 24.5, 25, 27.5, and 36 degree two-theta), TFF mannitol and leucine (e.g., 9.5, 20.2, 21, 24.5, 25, and 36 degree two-theta), and TFF pDNA formulation (e.g., 9.5, 20.2, 21, 24.5, 25, and 36 degree two-theta). The XRPD peak patterns of mannitol in these samples demonstrated that mannitol remained crystalline as a mixture of the δ and α forms [39]. Similarly, some peaks of leucine (e.g., 6 and 19 degree two-theta) were observed in the TFF neat leucine, TFF mannitol and leucine, and the TFF pDNA formulation, indicating that leucine remained crystalline after the process.

**Figure 4.**
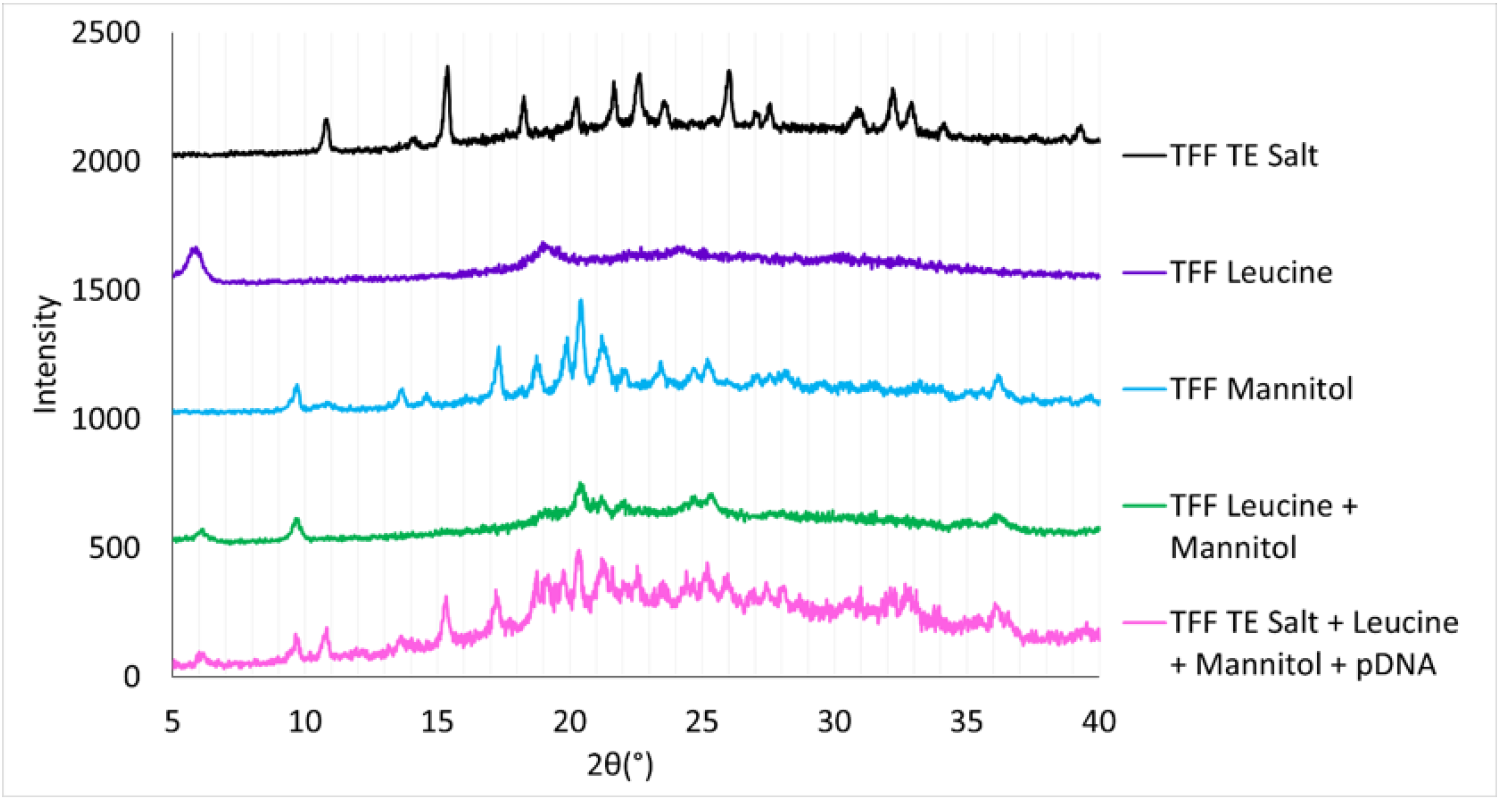
XRPD diffractograms of TFF pDNA formulation and excipients.

DSC was also used to determine the physical state of pDNA powder and excipients. The melting points of Tris and EDTA disodium salt were reported to be ∼175°C [40] and ∼242°C [41], respectively. Although XRPD diffractograms clearly demonstrated that Tris and EDTA disodium were crystalline after the process, no clear endothermic peak of Tris and EDTA disodium was observed on the DSC thermograms (Fig. 5, black line). Since both salts generally undergo thermal decomposition after melting [42], it is possible that the melting point of Tris and EDTA disodium was interfered by thermal decomposition. DSC thermograms showed that the melting point of TFF neat leucine and TFF neat mannitol was about 274°C and 167°C, respectively (purple and blue line). In the presence of mannitol, the melting point of leucine was decreased to 213°C (green line). Additionally, the melting point of leucine in TFF leucine and TE salt was further decreased to 197°C (yellow line), indicating that the presence of TE salt also contributed to the melting point depression of leucine. Although the presence of leucine did not result in the melting point depression of mannitol (∼166°C), the melting point of mannitol decreased to ∼135°C (orange line) when TE salt was combined with mannitol. Comparing DSC thermograms in the red and pink lines, the addition of pDNA in the formulation slightly decreased the melting point of mannitol and leucine (∼129°C and ∼216°C, respectively). No melting or glass transition temperature of pDNA was observed in the TFF pDNA formulation P7. Since the pDNA loading was only 5% in the formulation, the thermal events of pDNA were possibly below the detection limit of DSC analysis [43]. Finally, it is noted that both XRPD and DSC analyses were done using TFF pDNA powder formulation 7 that contained TE buffer. TFF pDNA powder formulation 3 that did not contain TE buffer was not analyzed, but data from our previously studies showed that TFF processed mannitol/leucine mixtures are crystalline as well [29, 44].

**Figure 5.**
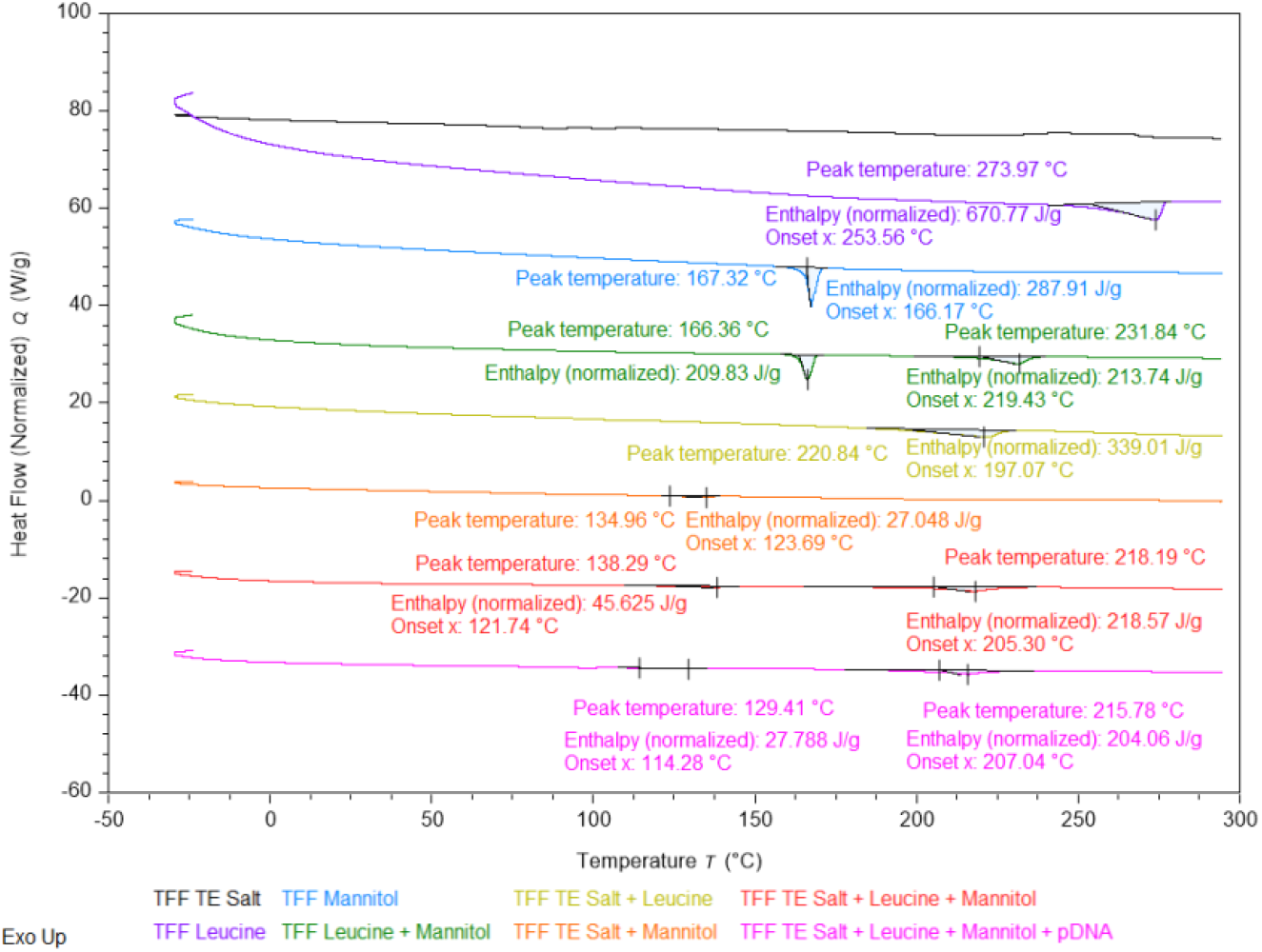
DSC thermograms of TFF pDNA formulation and excipients.

### Integrity of Plasmid DNA After Being Subject to Thin-Film Freezing

Plasmid powder formulation P7 had 5% pDNA loading, contained TE, and showed overall good aerosol performance properties. It was therefore chosen to test the integrity of the pDNA after it was subjected to TFF and reconstitution. As shown in Fig. 6, subjecting pCMV-β to TFF did not cause any significant change in the plasmid when comparing the plasmid before being subjected to TFF and reconstitution, with or without restriction digestion, in the agarose gel electrophoresis image, demonstrating that the TFF process did not compromise the chemical integrity of the plasmid.

**Figure 6.**
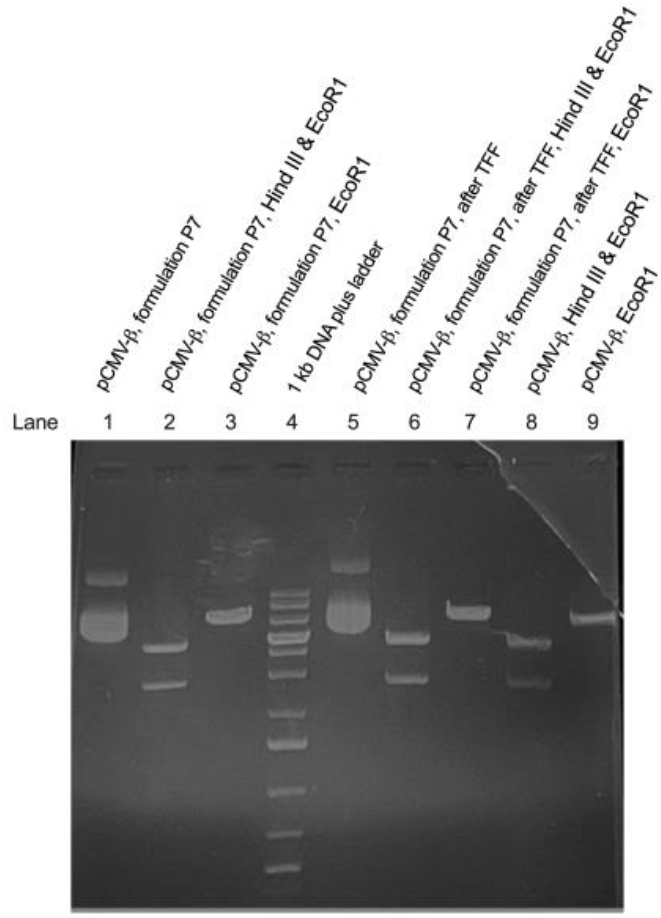
Chemical Integrity of pDNA before and after being subjected to TFF process. Lane 1, undigested pCMV-β in formulation P7 before TFF; Lane 2, pCMV-β in formulation P7 before TFF and digested with *Hind* III and *Eco*RI; Lane 3, pCMV-β in formulation P7 before TFF and digested with *Eco*RI; Lane 4, 1 kb plus DNA ladder; Lane 5, undigested pCMV-β reconstituted from formulation P7 powder; Lane 6, pCMV-β reconstituted from formulation P7 powder and then digested with *Hind* III and *Eco*RI; Lane 7, pCMV-β reconstituted from formulation P7 powder and then digested with *EcoRI*; Lane 8, original unformulated pCMV-β digested with *Hind* III and *Eco*RI; Lane 9, original unformulated pCMV-β digested with *Eco*RI. The loading of plasmid for Lane 1 and Lane 5 was 500 ng per well, other lanes were 420 ng.

### In Vitro Cell Transfection Study

To further investigate the integrity of the pDNA after being subjected to TFF, the pDNA’s ability transfect cells was tested in cells in culture. To do this, pVectOZ-GFP that expresses GFP protein was complexed with Lipofectamine-3000 to transfect A549 human lung epithelium cells. Agarose gel electrophoresis and restriction digestion with *Hind* III alone or *Hind* III and *Bam* H1 confirmed the integrity of the plasmid after it was subjected to TFF (data not shown). To optimize the pDNA dose in the *in vitro* cell transfection study, the effect of the pDNA dose on GFP expression in A549 cells was studied. As shown in Fig. 7A, overall, increasing plasmid dose led to higher levels of GFP expression; however, at the highest dose tested (i.e., 2000 ng/well), the GFP expression was lower than at 1000 ng/well. At all doses tested, no apparent cell toxicity was observed, likely because the dose of Lipofectamine was kept constant. The excess plasmids in the 2000 ng/well group may have contributed to the reduced GFP expression; when the ratio of the plasmid to the Lipofectamine was too higher, the resultant complexes may not be readily taken up by the cells. Nonetheless, the 500 ng/well dose was chosen to test the transfection efficiency of the pVectOZ-GFP before and after being subjected to TFF, and data in Fig. 7B showed that the TFF process did not significantly affect the activity of the plasmid, as there was not any difference in the GFP expression levels before and after TFF.

**Figure 7.**
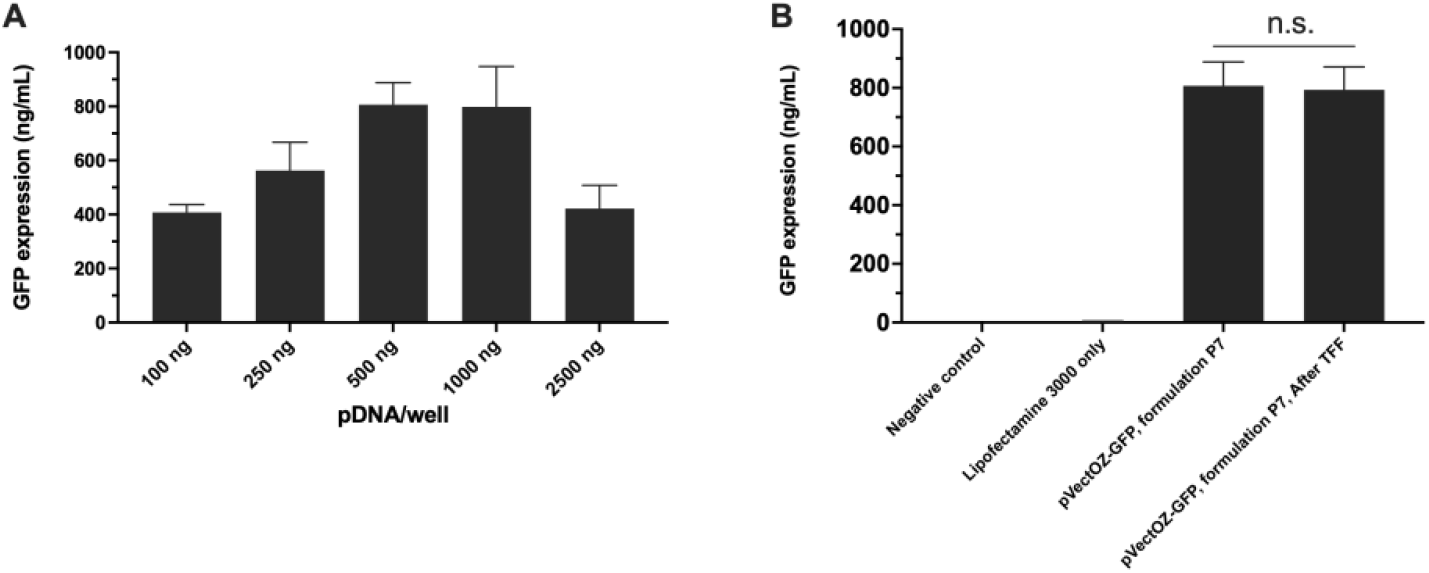
Cell transfection with pVectOZ-GFP before and after TFF. (A) Optimization of plasmid dose based on the GFP expression levels. (B) GFP expression in A549 cells after transfected with pVectOZ-GFP before and after it was subjected to TFF. Data are mean ± S.D. (*n* = 4) (n.s., not significant, t-test, two-tail, p > 0.05).

### Plasmid DNA Integrity After Actuation Using a Dry Powder Inhaler

The shear stress generated by nebulizing pDNA in liquid is known to cause damage to pDNA and can significantly reduce its transfection efficiency [8-10, 12]. Therefore, we investigated whether aerosolization of the TFF processed pDNA powder with a DPI device causes damages to pDNA. The pCMV-β plasmid was chosen in this study due to its larger size (i.e., 7.2k bp). We tested the integrity of pCMV-β after it was subjected to TFF and then aerosolized using a Plastiape NGI device. As positive controls, the pDNA in a solution that contained the identical excipients as in the powder was subjected to spray freezing or spray freeze-drying, as spraying is known to damage pDNA [13-16]. As shown in Fig. 8, no significant plasmid configuration change (e.g., nicking or linear form) was detectable after the plasmid was subjected to TFF (Lane 2 vs. Lane 3) and after the TFF pDNA powder was actuated from an DPI into NGI (Lane 2 vs. Lane 4), indicating that the plasmid would not be damaged when aerosolized as TFF powders using a DPI device into human lungs.

**Figure 8.**
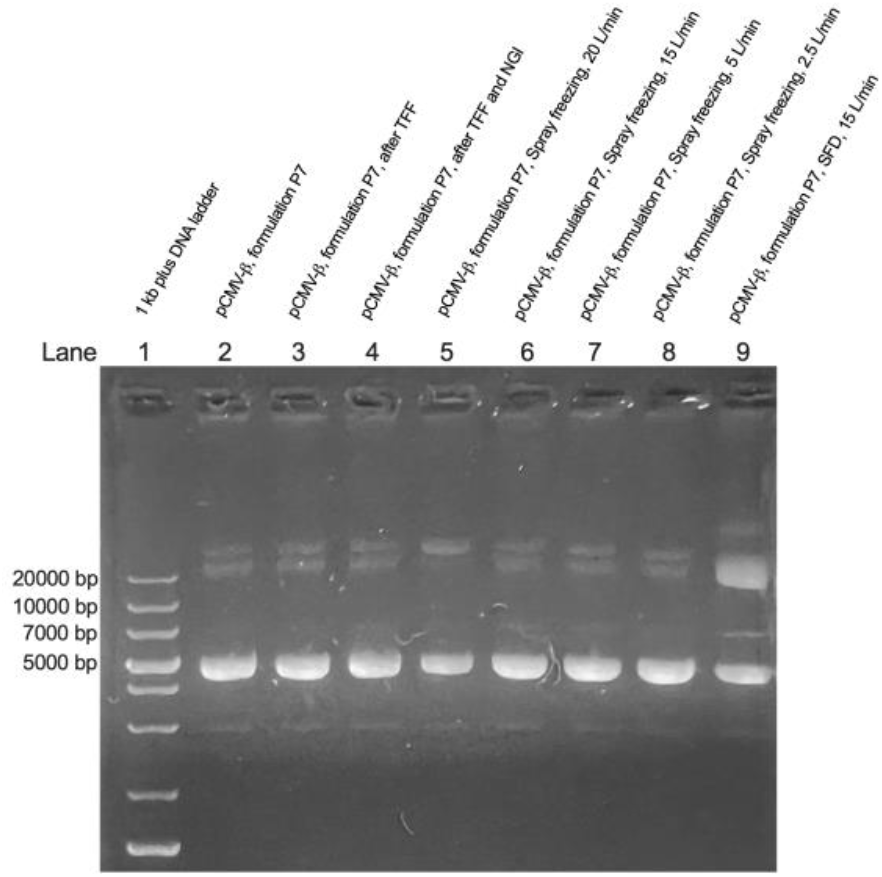
Plasmid DNA integrity after being subjected to TFF, TFF and actuation, spray freezing, and spray freeze-drying (SFD). Lane 1, 1kb plus DNA ladder; Lane 2, pCMV-β in formulation P7 before TFF; Lane 3, pCMV-β reconstituted from formulation P7 TFF powder; Lane 4, pCMV-β in formulation P7 TFF powder collected from NGI plates after the powder was actuated using a DPI; Lanes 5, 6, 7, and 8, pCMV-β in formulation P7 after spray freezing at air-flow rate of 20, 15, 5, and 2.4 L/min, respectively; Lane 9, pCMV-β in formulation P7 after SFD at air-flow rate of 15 L/min. The loading of plasmid was 500 ng per lane.

On the contrary, spray freezing at different air-flow rates (i.e., 20-2.5 L/min) caused changes in the plasmid (e.g., nicking as shown in Fig. 8, Lane 5, or various levels of linearization as shown in Lanes 6-8). The damages to the plasmid were likely from the shear stress during the spraying as pDNA is routinely subjected to freezing for longer term storage. Compared to spray freezing alone, spray freeze-drying showed a significant increase of linear and nicked forms of the plasmid (Fig. 8, Lanes 8 vs. 9), indicating the damages to the plasmid induced by the shear stress during the spray freezing step was amplified during the drying process. The finding is in agreement with reports by others showing the effect of shear stress during spray drying and spray freeze-drying on pDNA integrity. To overcome the damages caused the shearing, plasmids are often complexed with cationic polymers such as chitosan and polyethyleneimine (PEI), cationic liposomes or nanoparticles, or even encapsulated inside particulates such as poly lactic-co-glycolic acid microparticles [13, 15, 16, 45-47]. Unfortunately, those excipients and carriers are often associated with issues such as toxicities that limit their applications in humans [48, 49]. It is noted that cellular uptake of naked pDNA in its native form is relatively inefficient [50, 51], and complexing pDNA with cationic polymers or liposomes are usually employed to promote cellular uptake of pDNA [50, 52, 53]. Nonetheless, there is evidence that when naked pDNA is delivered into the lung of animals such as sheep, specific immune response against the antigens encoded by the plasmid can be induced [54, 55].

## CONCLUSION

It is feasible to apply TFF technology to engineer dry powders of pDNA with desirable aerosol performance, while preserving the chemical integrity and activity of the pDNA. In addition, the plasmid DNA in the dry powders is not sensitive to shearing when actuated using a dry powder inhaler.

## ACKNOWLEDGEMENT

Cui and Williams report financial support from TFF Pharmaceuticals, Inc.

